# A theory that predicts behaviors of disordered cytoskeletal networks

**DOI:** 10.1101/138537

**Authors:** Julio Belmonte, Maria Leptin, Nédélec François

**Affiliations:** Directors’s Research/Developmental Biology Unit, European Molecular Biology Laboratory, Meyerhofstrasse 1, 69117 Heidelberg, Germany; Cell Biology and Biophysics Unit, European Molecular Biology Laboratory, Meyerhofstrasse 1, 69117 Heidelberg, Germany

## Abstract

Morphogenesis in animal tissues is largely driven by tensions of actomyosin networks, generated by an active contractile process that can be reconstituted *in vitro*. Although the network components and their properties are known, the requirements for contractility are still poorly understood. Here, we describe a theory that predicts whether an isotropic network will contract, expand, or conserve its dimensions. This analytical theory correctly predicts the behavior of simulated networks consisting of filaments with varying combinations of connectors, and reveals conditions under which networks of rigid filaments are either contractile or expansile. Our results suggest that pulsatility is an intrinsic behavior of contractile networks if the filaments are not stable but turn over. The theory offers a unifying framework to think about mechanisms of contractions or expansion. It provides a foundation for the study of a broad range of processes involving cytoskeletal networks, and a basis for designing synthetic networks.

## Introduction

Networks of cytoskeletal filaments display a variety of behaviors. A decisive feature for the physiological role of networks is whether they contract or expand. For instance, actomyosin cortices can contract, and the tensions thus created determine the morphology of animal cells (Sanchez et al., 2011). Conversely, the mitotic spindle at anaphase is a network of microtubules that extends to segregate the chromosomes. Such behaviors are essential, but we still lack an intuitive understanding of how they come about, as it is difficult to extrapolate between the microscopic level, where filaments are moved by molecular motors and restrained by crosslinking elements, and the level of the entire system. Cytoskeletal filaments and many of their associated factors are well characterized biochemically. With sufficient knowledge of the relevant properties of the components of a particular network, it should be possible to predict the network behavior. Traditional approaches were particularly successful in predicting passive systems composed of reticulated polymers (Wolff and Kroy, 2012), and more recent developments in active gel theories address networks containing molecular motors (Prost et al., 2015). These latter theories however cannot explain the contractile or expansile nature of the network, as it arises from microscopic interactions that are not represented in the theory. To understand why contractility occurs, one must describe the system at higher resolution, and consider motors and filaments explicitly (Carlsson, 2006; Kruse and Jülicher, 2000; Lenz and Gardel, 2012; Liverpool et al., 2009; Liverpool and Marchetti, 2003; Ziebert et al., 2009). Small networks can also be studied with computer simulations (Bidone et al., 2017; Ennomani et al., 2016; Hiraiwa and Salbreux, 2016; Mendes Pinto et al., 2012; Oelz et al., 2015), but we miss a simpler approach that can make rapid predictions purely based on analytical deduction. Such a theoretical framework would be particularly valuable to classify the different behaviors that are seen experimentally. In search for such a general theory, we chose initially to concentrate on the major factor determining contraction of networks, that is the force created by molecular motors, although we recognized that filament shortening could also lead to contractility (Backouche et al., 2006; Mendes Pinto et al., 2012; Oelz et al., 2015). *In vitro* experiments have shown that contractility can arise with stabilized filaments. In such experiments, the filaments are initially distributed randomly, and molecular motors or crosslinkers added to the mixture make random connections between neighboring filaments. The active motions of molecular motors then drive network evolution. With microtubules and kinesin oligomers, static patterns such as asters (Köhler et al., 2011; Nedelec et al., 1997) or dynamic beating patterns (Katoh et al., 1998; Sanchez et al., 2011; Takiguchi, 1991; Thoresen et al., 2011) can arise. While radial (Backouche et al., 2006) and other patterns (Köhler et al., 2011) were also observed with actin, F-actin networks activated with myosin are predominantly contractile, as demonstrated in various geometries: bundles (Katoh et al., 1998; Takiguchi, 1991; Thoresen et al., 2011), rings (Reymann et al., 2012), planar networks (Murrell and Gardel, 2012), spherical cortices (Carvalho et al., 2013; Shah et al., 2014; Vogel et al., 2013) or 3D networks (Bendix et al., 2008; Koenderink et al., 2009). Microtubule networks with NCD or dynein motors are also contractile (Foster et al., 2015; Surrey et al., 2001). Several interesting mechanisms of contraction have been proposed and reviewed recently (Murrell et al., 2015; Salbreux et al., 2012), but each of these applies only to a particular system for which it explains the behavior. We propose here a theory that describes general principles of contractility and that can be applied to both microtubule and actin systems. We also illustrate that contractile systems become pulsatile if filament turnover is introduced in the model.

## Results

### A Simple Theory Predicts the Behavior of Random Networks

Let us consider a disorganized set of filaments connected by active and passive “connectors” made of two functional subunits (Fig. 1A, B). Possible passive connectors are crosslinkers such as Ase1, Plastin, alpha-Actinin or Filamin, whereas active connectors are oligomeric motors such as Myosin mini-filaments, Dynein complexes, bivalent motors such as Kinesin-5 or Myosin VI. By walking along filaments, bridging motors move the filaments relative to each other and change the network. It is however not obvious *a priori* how the sum of their local effects will influence the overall shape and size of the network. A computer can be used to simulate the dynamics of a network, but because all biochemical parameters must be specified in a simulation, only a finite set of conditions can be tested. We present here an analytical theory that overcomes this limitation. Active networks have been previously analyzed (Gowrishankar et al., 2012; Lenz, 2014; Liverpool and Marchetti, 2003; Nedelec et al., 1997; Ziebert et al., 2007) by considering pairs of filaments with one active connector (Fig. 1C). This approach is valid for sparsely connected networks in which only a few motors are active, but physiological networks must be well connected to exert force. In other words, the network should be elastically percolated, and there must exist continuous paths through which tension can be transmitted between any pair of distant points (Dasanayake et al., 2011). Specifically, we assumed that filaments are connected to at least two other filaments of the network. Focusing on one of these filaments (Fig. 1D), we see that the section of the filament between two connectors acts as a mechanical bridge between two points of the network. If the connectors are immobile, or if they both move in the same direction at the same speed, their distance remains constant, the section of the filament between them does not change in length, and the bridge is neutral (Fig. 1E). By contrast, if the two connectors move towards each other, the bridge exerts a contractile stress, whereas if they move apart, this produces an expansile stress (Fig. 1**E**).

**Figure 1:**
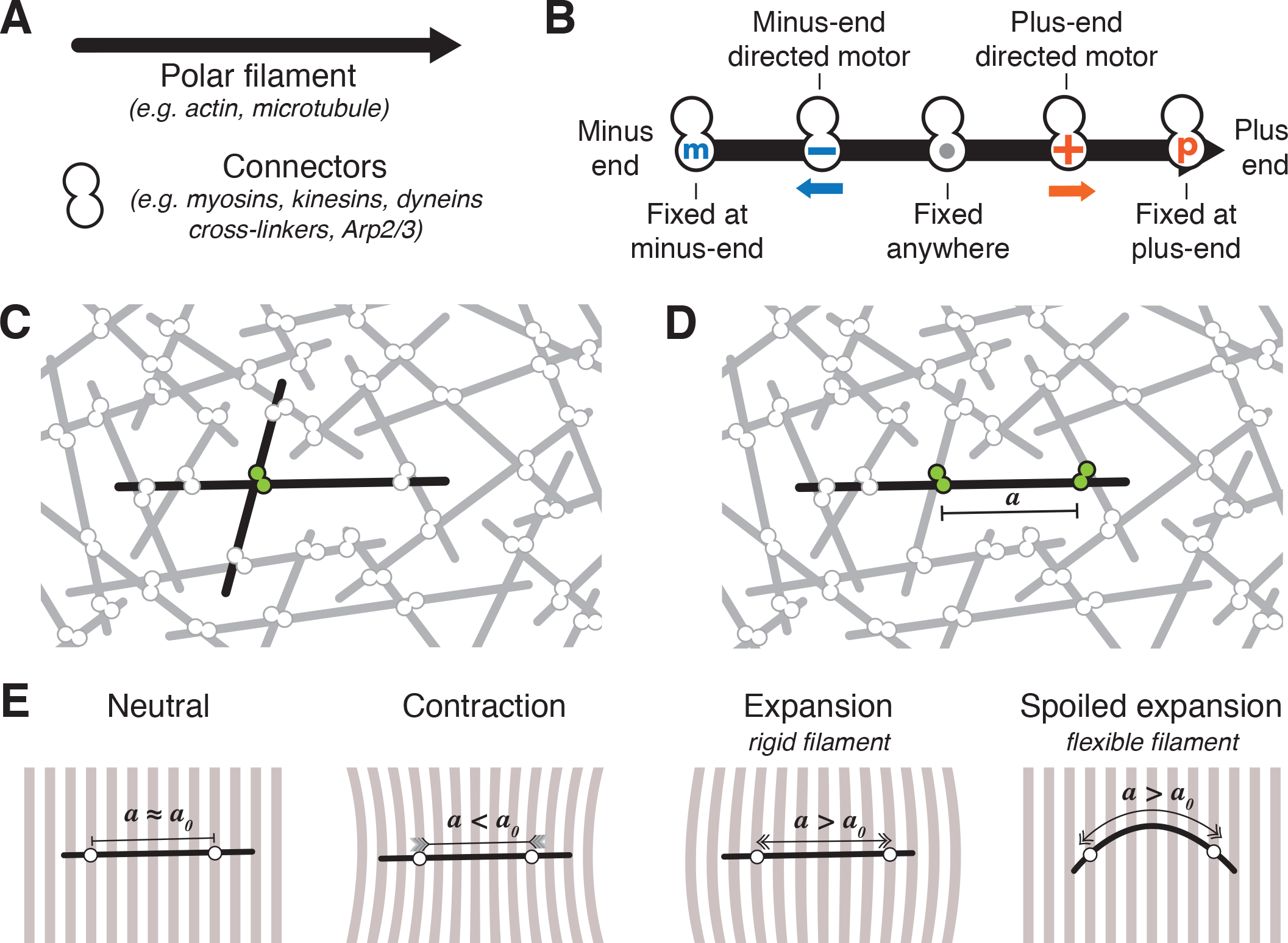
Elements of the theory. (**A**) Networks are composed of polar filaments that may bend, and connectors containing two subunits through which they can bridge two nearby filaments. (**B**) Subunits may be minus-end or plus-end directed motors that can bind anywhere to a filament, binders that can bind to any location along a filament, or end-binders that attach only at the minus or the plus ends of filaments. (**C, D**) To predict the behavior of a network, previous theories have considered a pair of filaments with a single connector between them, while the theory considered here is based on the effects that connectors bound to a sinle filament have on the rest of the network. (**E**) Pairs of connectors may generate local stress in the network depending on how the subunits move relative to one another on the filament. If the initial distance **a**_0_ between the subunits is maintained, the network does not deform. Local contraction is expected if the connectors move towards each other (**a**<**a**_0_) and expansion may occur if they move apart (**a**>**a**_0_). If the filamentis flexible, however, the expansile stress can be reduced if the filament buckles.

To predict if the whole network will contract or expand, we sum up the effects of all mechanical bridges in the network. To do this, we first list all the possible configurations for two connector subunits bound to a filament (shown below for two specific examples). For each configuration, we then calculate *da/dt*, where *a* is the distance between the two connectors measured along the filament. This quantity (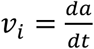 for configuration *i*) follows from the relative movements of the bound subunits of the connectors. When we sum all the contributions, we take into account the probability *p_i_* of each configuration to occur (∑*_i_ p_i_v_i_*), which can be calculated from the concentrations of components in the system, the binding and unbinding rate of the subunits, and other characteristics of the network (see Supplemental Information). We also distinguish the case where the filaments are rigid and can support expansile stress from the case where the filaments are flexible such that they buckle under compression. In the latter case, filament buckling spoils part or all of the expansile forces (Fig. 1E), and we thus discard the contribution of these expansile configurations. The summation of the weighted contributions of all configurations can then be carried out algebraically, and the sign of the result indicates the predicted network behavior.

### Actomyosin Networks with Motors and Crosslinkers

To develop the theory, we first applied it to a much studied model of cytoskeletal activity, that of actomyosin contraction, which has also been reconstituted *in vitro* (Carvalho et al., 2013; Katoh et al., 1998; Koenderink et al., 2009; Mizuno et al., 2007; Murrell and Gardel, 2012; Reymann et al., 2012; Shah et al., 2014; Takiguchi, 1991; Thoresen et al., 2011; Vogel et al., 2013). Actomyosin networks consist of stabilized F-actin filaments and two types of connectors: bivalent motors moving at speed *v* and passive crosslinkers (Fig. 2**A**). The crosslinker is composed of two identical subunits that may bind anywhere on the filaments, and that remain immobile until they detach. There are four possible ways to arrange the two types of connectors on a filament (Fig. 2**A**). Their likelihood depends on *P_M_* and *P_c_*, the probability of one or more motors and the probability of one or more crosslinkers being bound at an intersection of filaments, respectively. Two of the configurations are neutral in outcome: those with two motors and those with two crosslinkers. The other configurations are active and involve a motor and a crosslinker (Fig. 2**A**). There are two symmetric configurations with opposite outcomes. In one, the motor and the crosslinker approach each other at speed −*v*, and in the other they move apart at speed *v*. They have an equal likelihood that is proportional to *P_M_P_c_*(1 − *P_c_*), reflecting that one of the crossings should have at least one motor and no crosslinkers, with a likelihood *P_M_*(1 − *P_c_*), while the second crossing should have at least a crosslinker, with or without motor, carrying a likelihood *P_C_*. The net sum over the effects of all configurations in this example is null, and this predicts that a system made of rigid filaments that remain straight should neither contract nor expand. Contractile and expansile configurations cancel each other out, as found previously in the case where only motors were considered (Kruse and Jülicher, 2000). If the filament buckles, however, the expansile configuration will not be able to drive network expansion (Fig. 1E, last panel). Whether a filament buckles depends on the rigidity of the filament, the amount of force generated by the motors, and the distance *a* between the connectors. Under conditions in which the filaments always buckle, there are no expansile configurations, and the net sum is −*P_M_P_c_*(1 − *P_c_*)*v*. In this simple case, the sign reveals that the system is contractile, and this is correctly recapitulated in simulations, for example, when filaments are set to be very flexible (rigidity = 0.01 pN μm^2^) (Fig. 2BD and Movies 1 to 4).

**Figure 2:**
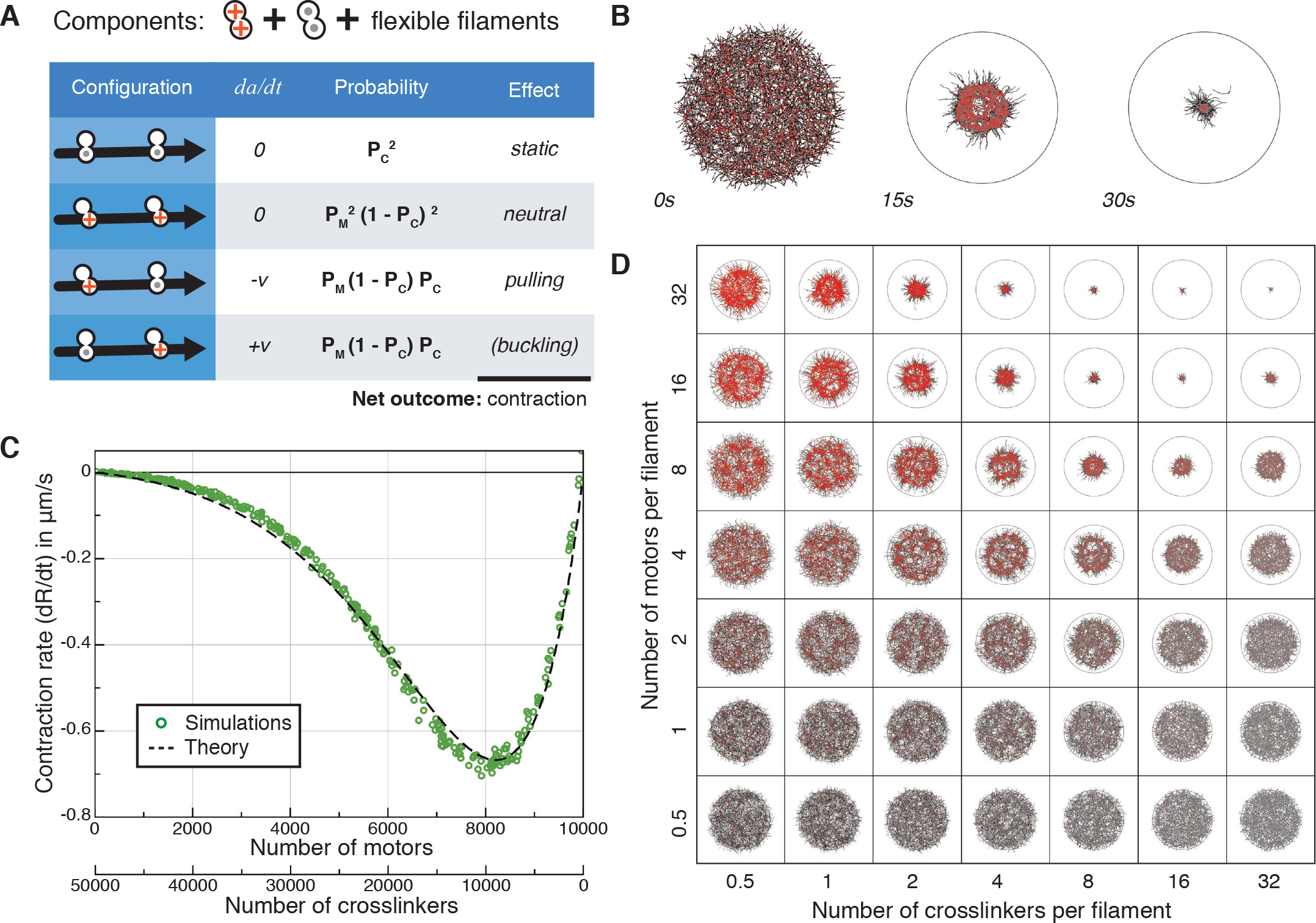
Predictions and simulations for actin-like networks of flexible filaments. (**A**) A system composed of flexible filaments and two types of connectors: crosslinkers and bivalent motors. The table lists the four possible configurations for two connectors bound to a filament, the relative movement of the connectors 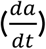, the likelihood, and the mechanical nature of each configuration. The likelihoods are combinations of *P_m_* and *P_c_*, *i.e*. the probabilities ofhaving a motor or a crosslinker at an intersection of two filaments (see Supplemental Information D3). (**B**) The evolution of a simulated random network composed of 1500 flexible filaments (bending rigidity = 0.01 pN μm^2^) and 12000 connectors of each type, distributed over a circular area of radius 15μm. (**C**) The contraction rate of a simulated network as a function of the ratio of crosslinkers to motors, with the total number of connectors kept constant. Each symbol indicates the result of one simulation. The broken line indicates the analytical prediction made by the theory (Supplemental Information D). (**D**) Snapshots at t=15s of networks similar to (**B**) containing varying numbers of motors (vertical axis) and crosslinkers (horizontal axis). No contraction occurs without crosslinkers or without motors, and the optimal contractile rate is obtained here for 8000 motors and 10000 crosslinkers. The location of the optimum can be predicted from the molecular properties of the connectors (Supplemental Information D).

### Networks of Semi-Flexible Filaments Contract at Predicted Rates

If the filaments are semi-flexible, which is the case for F-actin, with a rigidity of 0.075 pN μm^2^, the contribution of expansile configurations may not always be negligible, since a filament may or may not buckle depending on the length over which it is compressed. Therefore, to be able to predict the behavior of a network, it is necessary to know the conditions under which filaments buckle.

For an empirical assessment of this effect, we thus simulated networks in which the length and density of the filaments, and the number of crosslinkers and molecular motors were systematically varied. For this, we used Cytosim, an Open Source simulation engine that is based on Brownian Dynamics (Nedelec and Foethke, 2007). In brief, each filament is represented by a set of equidistant points, subject to bending elasticity (Fig. 3A). Crosslinkers and motors are represented by diffusing point-like particles, which bind stochastically to neighboring filaments (Fig. 3CD). Connectors with a stiffness *k* are formed when motors or crosslinkers are bound with their two subunits (Fig. 3E). The movement of motors follows a linear force-speed relationship (Fig. 3F). For simplicity, the unbinding rate is constant for this study, and a motor reaching the end of a filament immediately unbinds (Fig. 3G). Given a random network as initial condition, Cytosim simulates the movement of all the filaments in the system, and a contraction rate is extracted automatically (Supplemental Information C).

**Figure 3:**
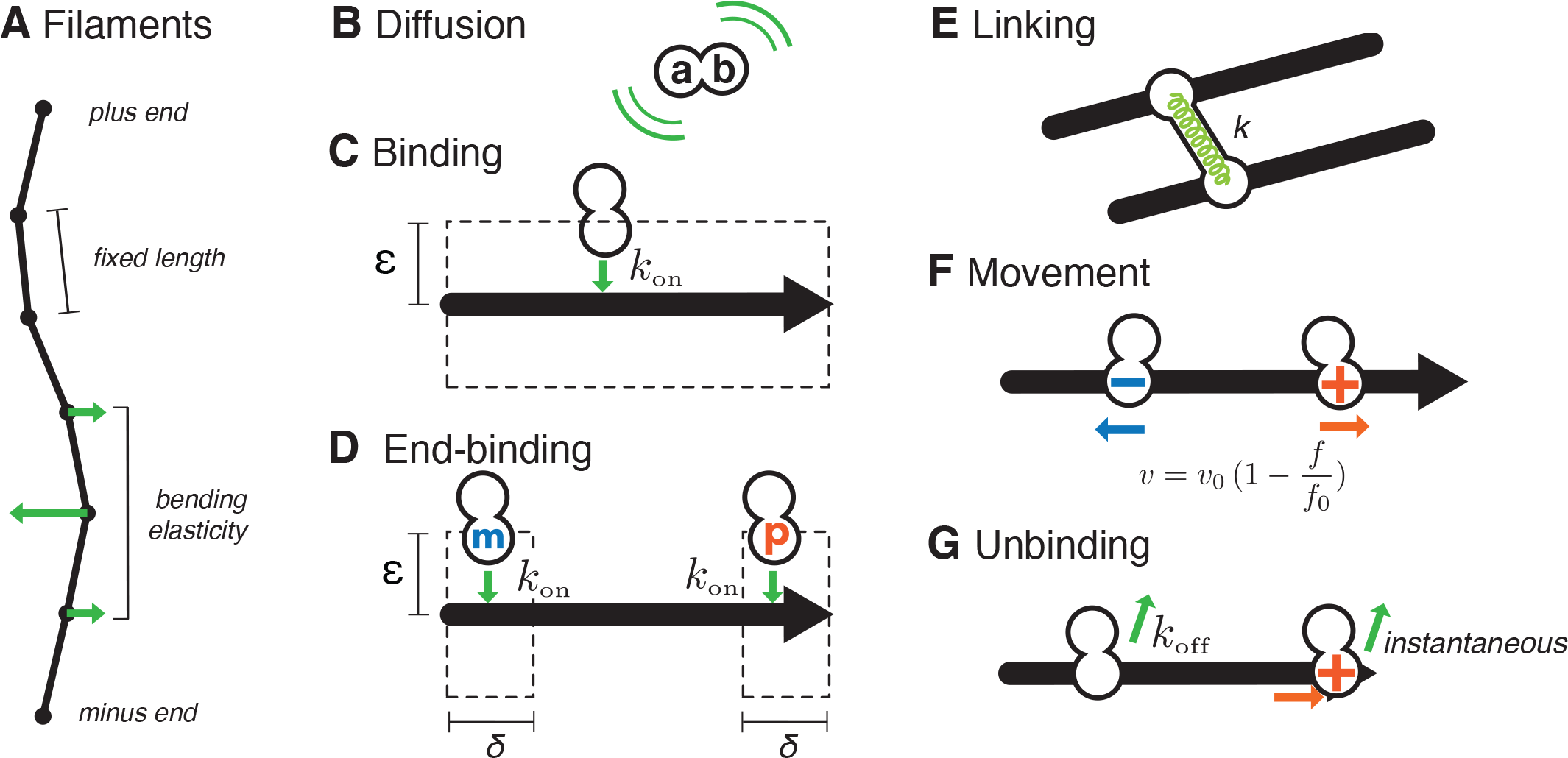
Elements of the stochastic model of cytoskeletal dynamics. (**A**) Networks of flexible filaments are simulated using a Brownian dynamics method. In brief, filaments are polar, thin and have a constant length. Each filament is modeled with an oriented string of points, defining segments of equal lengths. The movement of filament points follows Brownian dynamics, with elastic forces such as the bending elasticity of the filament, and the elasticity of connectors. (**B**) In the simulations, connecting molecules are made of two independent filament-binding subunits (a andb, which can be any one of those defined in Fig. 1B). When both subunits are unattached to filaments, the molecule diffuses within the simulation space. (**C**) Binding occurs at a constant rate k_on_ to any filament closer to than ε. Attachment occurs on the closest point of the filament. (**D**) End-binding follows the same rules as binding, but is restricted to a distance δ from the targeted filament end. (**E**) Connectors act mechanically as Hookean springs between two filaments, with stiffness k and zero resting length. (**F**) Motor subunits move towards either the plus- or minus-end of the filament with a linear force-velocity relationship. (**G**) All connector subunits detach with a force-independent rate k_off_, and motors detach immediately upon reaching a filament end.

Guided by the results of many simulations, we concluded that network contraction depends on the ratio between the mesh size *L*_1_ and the threshold distance *d*_0_ above which buckling occurs,which in turn can be calculated from the filament rigidity and the maximum force exerted by the motor (Supplemental Information D). If *L*_1_ < *d*_0_, then any filament segment of length *d*_0_ will be intersected by β=*d*_0_/*L*_1_ filaments. If any of these intersections is bridged by a crosslinker, this fixes the filament laterally and prevents it from buckling under the force of the motor(s) and crosslinker positioned at its ends. From these considerations, we can calculate the probability for a filament segment to buckle as 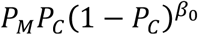 (Fig. 2C), where 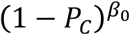 is the probability that none of the intersections between the motor and the crosslinkers are bridged by a crosslinker (Supplemental Information D). The theory predicts a dependence of the contraction rate on the number of connectors for a variety of conditions (Fig. S12–S14). For cases where the density of filaments is low, the contraction rate also depends on the contribution from a mechanism that has been anticipated long ago (Weisenberg and Cianci, 1984). In this configuration (Fig.7H), a motor acts on an intersection that is already connected by a crosslinker, producing a contractile force *by ‘zippering’* two filaments. For a network containing bivalent motors and bivalent crosslinkers, the dependence of contractility on the number of these connectors is thus predictable from first principles.

### Networks of Rigid Filaments Can Contract or Expand

We next explored systems composed of rigid filaments such as microtubules. Because some molecular motors are associated with microtubule ends in nature, we investigated the behavior induced by connectors comprised of motors and end-binding subunits (Fig. 4). As predicted by the theory (Fig. 4A), the simulations showed that the system is expansile if plus-end directed motors are combined with minus-end binding subunits (Fig. 4B, Movie 5), and contractile if plus-end directed motors are associated with plus-end binding subunits (Fig. 4C, Movie 6). A system composed of these two types of connectors can be either contractile or expansile depending on the relative concentrations of the connectors (Fig. 4DE, Movie 7). This reveals exciting avenues for the development of synthetic materials (Henkin et al., 2014) that could be tuned to be expansile or contractile. Using light-switchable molecular motors (Nakamura et al., 2014) would be particularly exciting in this respect.

**Figure 4:**
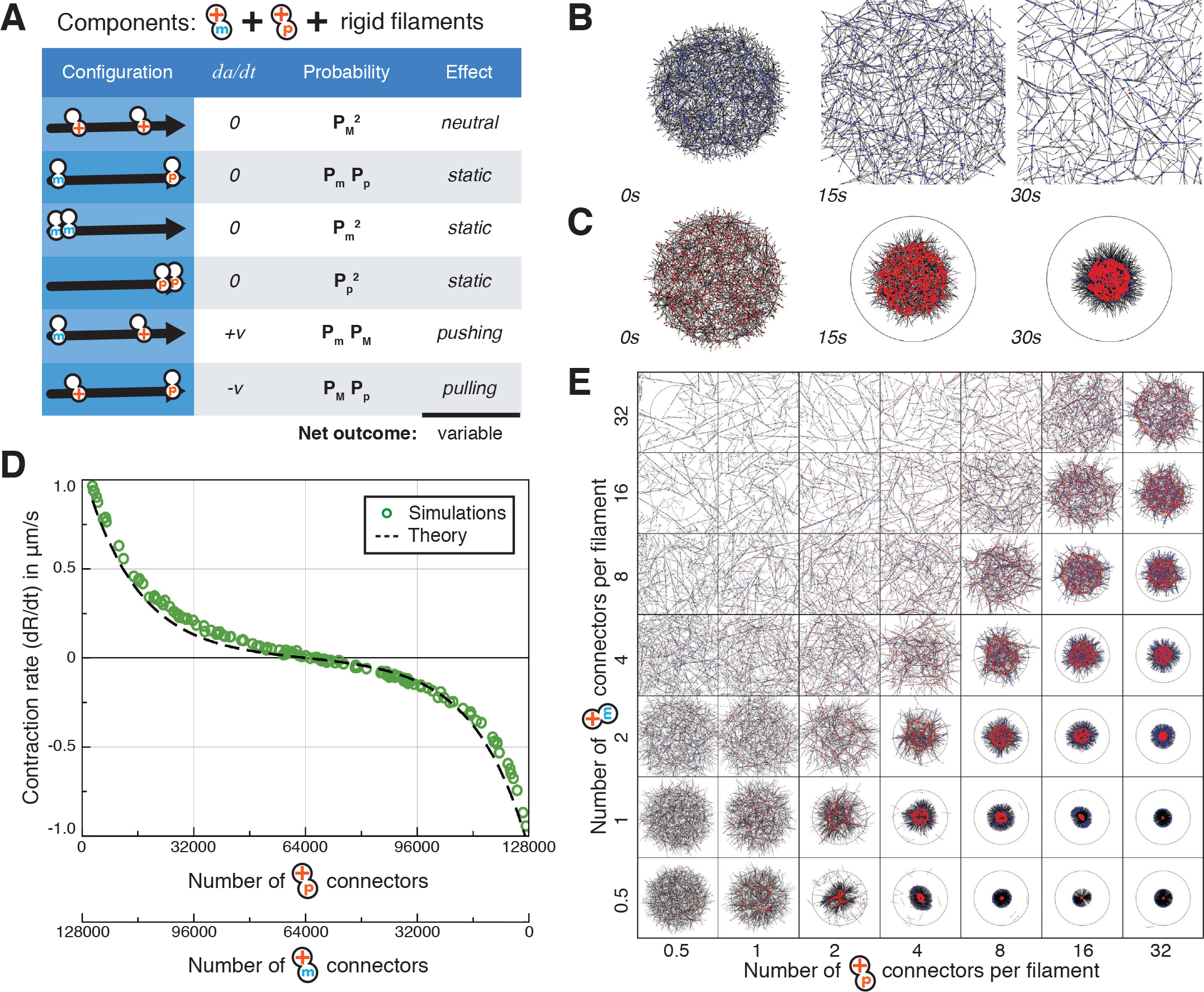
Predictions and simulations for microtubule-like networks of rigid filaments. (**A**) A system composed of rigid filaments and two types of connectors. One connector consists of a plus-end directed motor combined with a minus-end binder, the other is a plus-end directed motor combined with a plus-end binder. There are four possible configurations involving these two connectors. (**B**) Three time points on the evolution of an expansile network of 1500 straight filaments (their bendingrigidity is set as ‘infinite’ here) with 1500 motor-plus-end binders and 48000 motor-minus-end binders initially distributed over a circular area of radius 15μm. (**C**) Three time points on the evolution of a network similar as (**B**), but with 48000 motor-plus-end binders and 1500 motor-minus-end binders. (**D**) The contraction rate of a network as a function of the number of connectors, which are inversely varied. Each symbol represents a simulated random network of 4000 straight filaments initially distributed over a circular area of radius 25μm. Details of methods as in Fig. 2C. The broken line indicates the analytical prediction made by the theory (Supplemental Information F). (**E**) Simulations of networks containing varying numbers of connectors. Networks contain 1500 filaments initially distributed over a radius of 15μm. Depending on the concentrations of the connectors, the network can be expansile (top left corner) or contractile (bottom right corner). Snapshots at t=30s.

### The Effects of Many Combinations of Connectors is Predicted

To probe the general applicability of the theory, we simulated networks with mixtures of connectors containing 5 different types of subunits (Fig. 1B). A subunit can bind, and then either remain bound at the initial position, or move. Non-moving binders may be of a type that can bind anywhere on the filament, or they may be restricted in their binding to a region near the plus or the minus end. Moving elements (motors) can bind anywhere, but can be of two types, those moving to the plus and those moving to the minus end. By combining any two of these subunits, one can make 15 types of connectors. Simulated networks containing any one type of connector all behaved as predicted by the theory (see examples on Fig. 5A). We also simulated systems containing two different types of connectors (in equal quantities), both for flexible and rigid filaments. There are 210 possible combinations, and for every one of them, the simulations closely matched the prediction of the theory (Fig. 5BC, see Supplemental Information for details of the calculation). Many types of molecular elements that are found in nature, such as end-binding proteins, have not been used in reconstituted networks, but we can now predict what their effects on a network should be.

**Figure 5:**
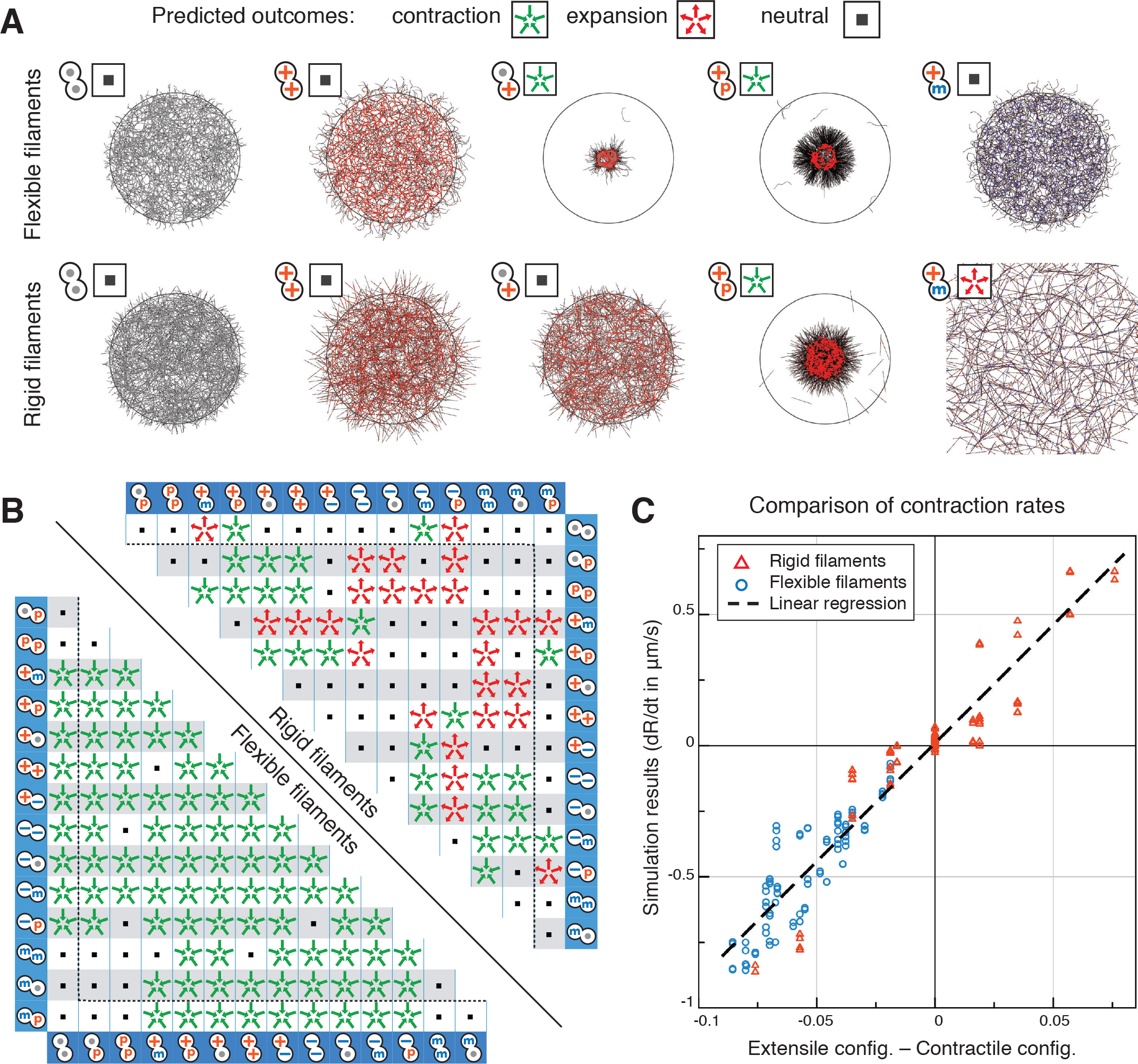
Additional predictions of the theory. The predictions of the theory for the various conditionsshown here are represented graphically as sets of green centripetal arrows for contraction, red centrifugal arrows for expansion, and grey squares for neutral networks. (**A**) Examples of simulations of networks with the indicated types of connectors. The predicted outcomes of network contraction, expansion or neutrality (symbol at the top left of each simulation) are confirmed by the behaviorof the network in simulations in each case. The networks are composed of 1500 flexible or rigid filaments, and 24000 connectors. Snapshots at t=20s. (**B**) Summary of the predictions for random networks with all possible combinations of all possible types of connectors, either with flexible (bending rigidity = 0.01 pN μm^2^, below diagonal) or rigid filaments (straight filaments, above diagonal). The networks contain 4000 filaments and 64000 connectors at a 1:1 ratio. The labels of the rows and columns of the table indicate the type of the two connectors that are mixed. (**C**) Comparison of the contraction rates predicted by the theory (horizontal axis) with the rates obtained by simulation (vertical axis). Each data point indicates one of the 210 systems considered in (**B**). Networks are made of 4000 filaments and 64000 connectors initially distributed over a circular area of radius 25μm. In this case, all the binding parameters of the subunits and the concentration of connectors are always equal, such that the prediction is simplified (Supplemental Information E).

### Heterogeneous systems composed of different types of filaments

So far, we have considered systems made of one type of filament, but some networks *in vivo* contain different types of filaments. In particular myosin II motors are organized into thick antiparallel ‘mini-filaments’ with an average length of 300 nm (Verkhovsky and Borisy, 1993). Networks such as the actomyosin meshwork of the cell cortex and the contractile actin cables in cells are thus heterogeneous systems in which F-actin filaments are mixed with mini-filaments, which are also the motors driving the system out of equilibrium. To probe if the theory could hold for such heterogeneous systems, we listed all the possible combinations of two connectors for the two types of filaments (Fig. 6A). Similar to the homogeneous case (Fig. 2) this analysis predicts that the system should be contractile if crosslinkers are also present, and neutral otherwise. We then simulated such a system of actin-like filaments and mini-filaments composed of a rigid backbone of length 0.5μm with a motor subunit at each end. The results confirmed the predicted behaviors (Fig. 6B and 6C), suggesting that the theory can be applied to heterogeneous networks. It is tempting to think that the approach can also be extended to anisotropic networks if the probabilities of the configurations are calculated locally.

**Figure 6:**
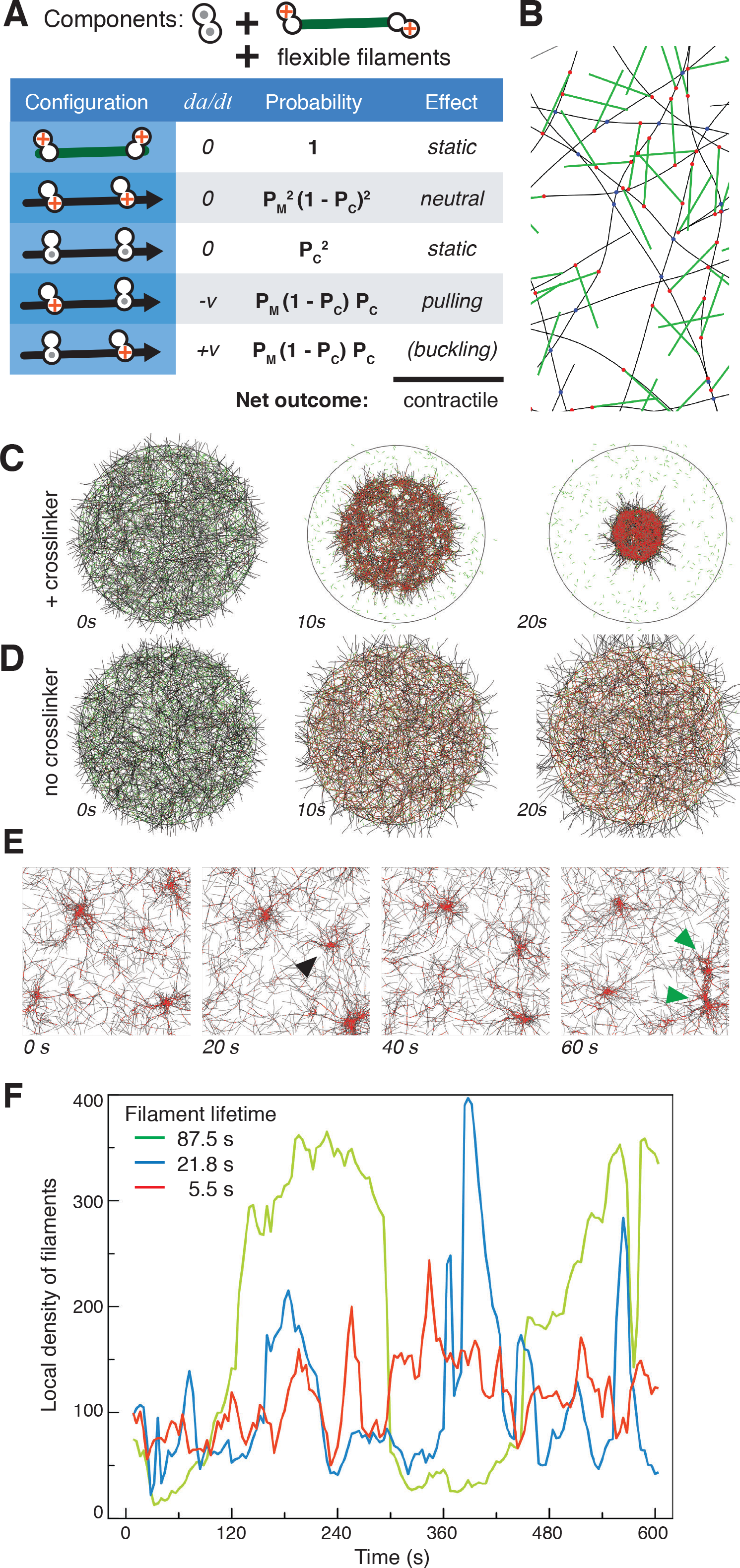
Heterogeneous and pulsatile systems. (A) Configurations present in a heterogeneous network containing rigid minifilaments and flexible actin-like filaments. The motors are permanently attached at the extremities of the minifilaments, such as to represent myosin minifilaments. The system is predicted to be contractile in the presence of passive crosslinkers connecting actin filaments directly, and neutral without crosslinkers. (B) Detail of a simulation with minifilaments (green) and crosslinkers (blue). (C, D) The simulated systems contract with crosslinkers, but not when they are omitted. (E) Time series of a simulation with filament turnover, 1400 filaments (rigidity 0.075 pN μm^2^), 22400 motors, 5600 crosslinkers within periodic boundary conditions with size 16 μm. Filament turnover was implemented by deleting a randomly selected filament and placing a new filament at a random location, stochastically with a rate R=64s^−1^, corresponding to an average filament lifetime of ~21.8s. The series shows the formation of a new contractile spot at the right side (black arrowhead) and its downward movement and fusion with the contractile spot at the lower right corner of the panels (green arrowheads). (F) The local density of filaments in an arbitrarily chosen region covering ~6% of the simulated space as a function of time. The data with filament lifetime 21.8s are from the simulation shown in (A). The network continues to redistribute, showing irregular variations of the local filament density, and does not contract into one spot.

### Contractile Systems Pulse if Filaments Turn Over

So far, we have considered systems made of filaments of fixed length that persist infinitely. Under these conditions,network contraction and expansion are non-reversible events, that only occur once. This is indeed what happens with most *in vitro* reconstituted cytoskeletal networks, obtained with stabilized filaments (Carvalho et al., 2013; Foster et al., 2015; Katoh et al., 1998; Murrell and Gardel, 2012; Shah et al., 2014; Surrey et al., 2001; Takiguchi, 1991; Thoresen et al., 2011; Vogel et al., 2013). But how does this relate to networks *in vivo*, which manage to avoid such a collapse? The simulations described above do not perfectly mimic the situation *in vivo*, since cytoskeletal filaments elongate by self-assembly and if they remain dynamic will eventually vanish, such that both the length of the filaments, and their abundance are fluctuating quantities that can be regulated. In some cells,actomyosin networks persist for several minutes, and exhibit pulsatile behavior that is reminiscent of contractility (He et al., 2010; Martin et al., 2009; Munro et al., 2004; Solon et al., 2009). To test the relationship between this dynamic behavior and contractility, we extended our simulations to include filament turnover. Specifically, to implement an average lifetime T for the filament, we randomly selected and deleted one of the N filaments at a rate N/T, and replaced it with a new one placed at a random position (Fig. 6D). We typically observed that within 3s < *T* < 200s, a configuration that was contractile without turnover now displayed pulsed contractions (Fig. 6E, Movie 8). Note that these new simulations used periodic boundary conditions (PBC), because if the system is allowed to contract freely, pulses cannot be observed. Use of PBC imposes a constant surface on the system and thereby forces the network to build up tension. It corresponds best to a network that is attached at the cell boundaries, without requiring additional assumptions on the nature of the attachment. These results confirm earlier models that considered filament dynamics (Bidone et al., 2017) or turnover (Hiraiwa and Salbreux, 2016; Kumar et al., 2014), illustrating that with filament turnover, a system that was contractile otherwise can be pulsatile. As suggested by a coarse-grained model (Kumar et al., 2014), we wondered if pulsatility was a general consequence of turnover. We thus simulated networks with all the combinations of connectors that were contractile on Fig.5B and varied systematically the filament turnover rate. All displayed pulsatile behavior, confirming the universal nature of the phenomenon (data not shown).

## Discussion

The theory we present here predicts the initial evolution of a network from the properties of its connectors. We have confirmed these predictions with simulations for all tested conditions. The model implemented in the simulation is intentionally minimalistic, as subunit binding, unbinding and filament turnover occurred at constant rates and independently of other events. All simulations where done in 2D, and did not consider steric interactions between the filaments, which in 2D would induce artifacts. We expect our theoretical arguments to hold also for other types of networks such as filament bundles or 3D networks. However, the calculation presented in the supplemental information depends on the geometry of the network, and would need to be revised to apply to these different conditions. Our analytical prediction of network behavior was based on the characteristics of the connectors but did not include physical parameters such as the viscosity of the medium. Rather than an absolute contraction rate, this theory predicts the relative contraction rates of two networks when the parameters of the connectors (numbers, types, binding rates, unbinding rates) are different between them. Such a prediction is immediately valuable, as it can be readily tested experimentally by systematically varying the concentration of both motors and crosslinkers in reconstituted *in vitro* networks.

For a system containing crosslinkers and bivalent motors (Fig. 2), our analysis indicates that the ‘active’ contractile configurations involve both a crosslinker and a motor. We thus expect these two types of activities always to be found in a contractile system. However, we note that in an *in vitro* experimental system, the assumedly pure preparation of motors that is added may in fact contain damaged ‘dead’ motor proteins that act as passive connectors. Nevertheless, addition of crosslinkers indeed dramatically enhances the effect of myosin, a phenomenon observed more than 50 years ago (Ebashi and Ebashi, 1964). For 2D networks (Fig. 2CD and Supplemental Information), we could explain why the maximum contractility is obtained *in vitro* with approximately equal amounts of motors and crosslinkers (Bendix et al., 2008; Ennomani et al., 2016; Köhler and Bausch, 2012). Our theory also explains that under the action of myosin VI, a branched network made with Arp2/3, which represents an example of a connector combining end-binding and side-binding activities together, is more contractile than a network connected by crosslinkers binding anywhere along the filaments (Ennomani et al., 2016). This is because, as myosin VI is directed to the pointed end, the configuration involving a crosslinker bound to the pointed end (Arp2/3) is always contractile. Thus,at equal levels of connectivity, a network made with Arp2/3 is more contractile than a network made with a non-specific crosslinker, and less contractile than a network made with only end-to-end crosslinkers.

Because we focused on the initial behavior of a network, it seemed justified to ignore mechanisms that regulate the characteristics of the cell cortex, for instance its thickness. We used however simulations to explore the effects of filament turnover (Fig. 6DE). The fact that introducing filament turnover was sufficient to induce pulsing in all the contractile scenarios leads to the surprising conclusion that pulsatility is an intrinsic behavior of contractile networks of non-stable filaments, and that no other external elements are necessary. This of course does not mean that in the natural biological situation there may not be regulatory elements superimposed on the underlying mechanism that suppress or enhance pulsing (Nishikawa et al., 2017). Pulsing is seen only over a certain range of filament lifetimes (Fig. 6D), indicating that one such regulatory input could be via the stability of filaments: for example, stabilizing filaments should, according to our simulations, arrest pulsing, and the ability of myosin to destroy filaments (Matsui et al., 2011) could on the contrary enhance pulsing. The characteristics of the pulses can be further tuned by modulating the other characteristics of the network components, as has also been seen *in vivo* for the regulation of myosin (Munjal et al., 2015). Pulses may be a desired feature or an inevitable consequence of having filament turnover when the connectors lead to contractility. This important feature of *in vivo* cytoskeletal networks deserves more investigation in the future.

Importantly, our theory unifies ideas previously proposed for various biological systems, and we will discuss now how various mechanisms giving rise to contraction or extension fit within the new theory (Fig. 7).

**Figure 7:**
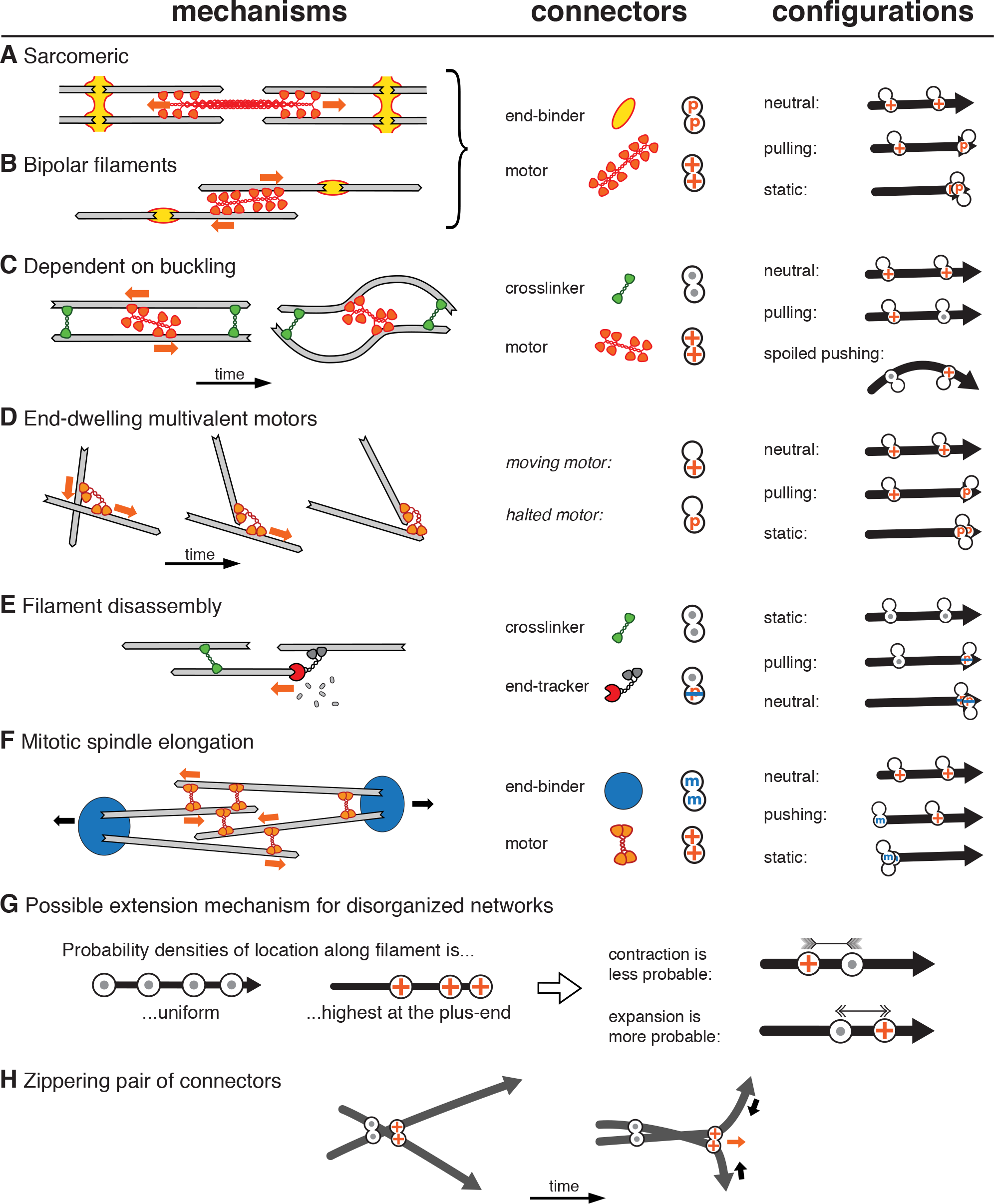
Review of contractile and expansile mechanisms. Previously described mechanisms can be considered by focusing on pairs of connectors present on filaments. The sarcomeric (A) and the analogous mechanism involving bipolar filaments (B) have a barbed-end directed multivalent motor acting on filaments that are connected at their pointed ends by molecular complexes that act as connectors. These systems are always contractile because there are only two active configurations: one involving two motors, which is neutral, and one with a motor and an end-binder, which is contractile since the motor always moves towards the end binder. The buckling-dependent mechanism (C) leads to contractility because the flexibility of the filament spoils the expansile configuration. Thus, if the filaments are sufficiently flexible, the net effect will be contractile (see Fig. 2). In systems containing only end-dwelling multivalent motors (D), the motors generate contraction without added passiveconnectors, because they eventually come to a halt at the end of the filament and thereby act as end-binding connectors. Configurations involving a motor halted at the end, and a motor moving towards this end along the same filament result in contraction. Note that since in this mechanism no configuration is expansile, and the net effect is always contractile, irrespective of filament buckling. A connector with a subunit that binds to a disassembling end of a filament (**E**) generates only one activeconfiguration which is always contractile, even in the absence of motors. In this example, the end-tracker binds to the barbed end, and moves towards the pointed end by tracking a depolymerizing end (or inducing its disassembly). (**F**) A mitotic spindle at anaphase may be considered as a network held together by multivalent plus-end directed motors from the Kinesin5 family, and by factors connecting the microtubule minus ends at the spindle pole. With these two types of connectors, the configurations involving two connectors are neutral, passive, or expansile. (**G**) A system can be made expansile by the “antenna effect”, because motors acquire an asymmetric distribution profile along the filaments. In the presence of this effect, contractile configurations are less likely than expansile ones, and the overall system can become expansile as a consequence. (**H**) Some mechanisms of contraction involve two connectors acting on more than one filaments. In the casedepicted here, two crossing filaments will be “zipperred together”, by a pair of connectors moving apart. This configuration is able to create a contractile force dipole in the direction perpendicular to the filaments.

In the sarcomeres found in striated muscles, myosin II mini-filaments pull on filaments arranged in an anti-parallel manner (Fig. 7A). This system can be seen as containing two types of connectors: a passive one linking the barbed ends of the filament, and a motor directed towards the barbed end. Three possible combinations can be made with these two connectors (Fig. 7, right column). In the absence of any expansile configuration, the system is bound to always be contractile. A sarcomeric system is highly ordered,but less organized systems made of the same subunits, for example bipolar filaments that are discussed for smooth muscles (Fig. 7B) or a fully disorganized system (Fig.5A, fourth column) are also contractile.

For a system in which the crosslinkers are not restricted to binding to the filament ends, but can bind anywhere along the length (Fig. 7C) both contractile and expansile configurations arise. Following the discussion on how buckling promotes contraction of a disorganized actin network (Liverpool et al., 2009; Mizuno et al., 2007) we argued that buckling can spoil some of the expansile configurations, tipping the balance in favor of contraction.

One popular mechanism to explain the contraction of microtubule networks (Fig. 7D) does not require filament bending, but involves a motor that has an affinity for the end of the filaments (Hyman and Karsenti, 1996; Nedelec et al., 1997). Because the motor walks towards the end, where it may be transiently trapped, configurations are contractile or neutral, but never expansile, and the entire network itself is therefore contractile (Foster et al., 2015). Looking at the set of configurations (Fig. 7, right column), the similarity of this mechanism with sarcomeric contractility (Fig. 7AB) is noticeable. In the case of the end-dwelling motor, however, the same molecular type is involved in generating the active and neutral end-binding connections.

Although we did not consider filament disassembly in this study, we would suggest that the theory can be applied also to this situation. For example, a molecule that tracks and remains bound to the depolymerizing end of a filament (Fig. 7E) will reduce the distance between itself and a connector located elsewhere on the filament, thereby creating a pulling force. By calculating the likelihood of such a configuration, one may be able to predict the overall contractility of the network. We also did not consider filament elongation, which is a prominent mechanism by which actin networks expand, but identify conditions where, even without this mechanism, a network can increase its surface area.

A system with expansile configurations will extend if the filaments are sufficiently rigid to resist buckling, which is more likely to be the case for microtubules than for actin. A mitotic spindle is a complicated structure, which can be found in multiple states. In most cells metaphase corresponds to a steady state in which contractile contributions must on average compensate expansile ones. In *Xenopus laevis*, contraction is driven by the motor minus-end directed Dynein (Foster et al., 2015) while expansion is driven by the plus-end motor Kinesin5 (Needleman and Brugués, 2014). When anaphase is induced, this balance is broken leading to spindle elongation, and chromosome segregation. For the sake of the argument, we consider here that the spindle at anaphase is purely driven by kinesin5, and thus assume only two types of connectors (Fig. 7F): passive complexes containing the protein NuMA, which connect the minus-ends of microtubules at the spindle poles, and plus-end directed motors connecting adjacent, antiparallel microtubules. Since Kinesin5 walks away from the minus-ends, the configurations are symmetric to the sarcomeric systems (Fig. 7A), and expand the anaphase spindle. A disorganized network made of the same connectors is also expansile (Fig. 4**B**).

Additional expansile microtubule systems can be found in blood platelets and their progenitor cells, the megakaryocytes. During pro-platelet generation, the microtubules assemble into bundles that elongate, and this elongation is dependent on the activity of the molecular motors Dynein (Patel, 2005). In the mature platelets, microtubules are organized into a closed circular bundle, and this bundle must be able to resist contractile forces as it pushes outward on the plasma membrane. It was recently reported that the microtubule ring elongates after platelet activation, in a manner that is dependent on microtubules motors, but we lacked until now a microscopic picture of the elongation mechanism (Diagouraga et al., 2014). Our systematic exploration of random networks (Fig. 5) suggests different scenarios that could explain why the system is expansile. Beyond the relevance to these *in vivo* systems, it will be exciting to follow these principles to create expansile networks of microtubules *in vitro*, since end-binders are available to synthetic biologists.

Finally, in a system where the symmetry provides an equal number of contractile and expansile configurations, any imbalance in the probabilities of these configurations may lead to overall contraction or expansion (Gao et al., 2015). Following this principle, we can suggest here an explanation for the expansile nature of *in vitro* networks (Sanchez et al., 2012). Particularly, if the motors are sufficiently processive, they may run a distance that is comparable to the length of the filament, and in this case their distribution along the length of the filament will be non-uniform (Fig. 7G). This effect has been called the antenna effect (Varga et al., 2006), and arises as a consequence of the motility, in a situation where binding has the same probability at every position of the filament. A plus-end directed motor would accumulate towards the plus ends of microtubules (Fig. 7G). In the presence of crosslinkers, such an effect will increase the likelihood of the expansile configurations, and lower the likelihood of the contractile configurations, thus promoting expansion. It is interesting to note that even if the motors were directed towards the minus-end of microtubules, the antenna effect would still lead to a bias in favor of expansion. The net imbalance will depend on the biophysical properties of the motors (speed, unbinding rate), and the length of the microtubules, and could provide tunable expansibility for networks made *in vivo* with microtubules (Sanchez et al., 2012).

In conclusion, our theory offers a framework for elementary mechanisms of expansion or contraction. It is a starting point for further exploration, since in its current state, the theory does not explain all the phenomena observed in simulations. In particular, for very flexible filaments a configuration involving two connectors starting from the same point of contact between two filaments seems to contribute to contraction (Fig. 7H). In this case, the connectors can pull the network together, even though the distance between them is growing. This interesting effect, which is analogous to a zipper, can only be understood by considering two filaments and two connectors, whereas our theory considered one filament and two connectors. Because this effect only contributes little in most cases (see Supplemental Information D5), the theory is still able to make accurate predictions, even for complex systems (Fig.5BC). Mathematically, the theory could be understood as providing a first-order approximation of the exact contractile rate.

From the theoretical framework presented here, with its clear predictions, perhaps a classification of the different types of active networks found in nature will emerge. Our approach may also inspire novel avenues for synthetic filament networks with enhanced functionalities.

## Author contributions

This work arose from regular discussions between the authors, which all have contributed significantly to the findings. All authors together wrote the article.

## Acknowledgments

We are thankful to EMBO and EMBL for support, in particular for its high-performance computing services. We thank members of the Leptin and Nedelec groups, particularly H. Turlier, S. Dmitrieff, M. Lera Ramirez and Y. Jeske, as well as S. Blandin, D. Gilmour, T. Hiiragi, J.-P. Shen, R. Prevedel and P. Lenart for critically reading the manuscript. J.M.B. is the recipient of an EMBL Interdisciplinary Postdoctoral fellowship, which is co-funded by Marie Curie Actions of the European Commission.

